# The SARS-CoV-2 nonstructural protein 15 collapses the host cytoskeleton by interacting with the host keratin type II cytoskeletal 1

**DOI:** 10.64898/2025.12.25.696538

**Authors:** Donald Tam, Cesar Monjaras-Avila, Hans Adomat, Horacio Bach

## Abstract

COVID-19, the deadliest recorded pandemic of the 21^st^ century, was caused by a novel coronavirus, SARS-CoV-2. To explore why this coronavirus was significantly more pathogenic than its predecessors, more information is needed about the functions of the viral proteins that contributed to this uptick in virulence. This study explored how one of these viral proteins, the nonstructural protein 15 (NSP15), interacts with the host. Our findings revealed a novel protein-protein interaction between NSP15 and the host keratin type II cytoskeletal 1 in Vero E6 cells, as determined by mass spectrometry, and subsequently validated by immunoprecipitation. We observed that when NSP15 was delivered into Vero E6 cells, the fibrous network of keratin intermediate filaments was disrupted, and the nuclear structure was lost. This disruption of the keratin cytoskeleton was then shown to cause a statistically significant reduction in cell viability of Vero E6 cells. These results indicate that the presence of NSP15 in host cells leads to a collapse of the keratin cytoskeleton. This cytoskeletal collapse could be a mechanism the virus employs to escape the infected host cell via cell lysis, once significant amounts of viral protein and progeny have accumulated.

## Introduction

In the past two centuries, there have been four pandemics caused by various influenza strains, including H2N2 (1889 and 1957), H1N1 (1918), and H3N2 (1968) [1]. However, in recent years, other pandemics caused by coronaviruses have risen, with the Severe Acute Respiratory Syndrome (SARS) outbreak in 2001 followed by the Middle East Respiratory Syndrome-related Coronavirus (MERS-CoV) outbreak in 2012. The most recent pandemic to date was the Coronavirus Disease 19 (COVID-19) pandemic, marking the 3^rd^ coronavirus epidemic in under 20 years.

COVID-19 is a respiratory disease caused by the Severe Acute Respiratory Syndrome Coronavirus 2 (SARS-CoV-2) reported in Wuhan, China, in 2019. This virus quickly spread worldwide and became a pandemic. This pandemic caught the world by surprise, leading to significant lockdowns worldwide that uprooted the global economy and claimed millions of lives. Although the pandemic has now concluded, its lasting impacts highlight the importance of preparation for the next pandemic. As such, actively exploring and understanding the factors that contributed to the high pathogenicity and survival of SARS-CoV-2 and other coronaviruses will be essential for guiding the development of better treatments to target this family of viruses in future outbreaks.

SARS-CoV-2 is an enveloped β-coronavirus with a single-stranded, positive-sense, with a post-transcriptionally modified (5’cap and 3’poly-A tail) RNA genome that spans 29.9 kilobases [2,3]. The SARS-CoV-2 genome encodes three classes of proteins: structural proteins, nonstructural proteins (NSPs), and accessory proteins. The first class consists of four structural proteins: the envelope protein (E), the membrane protein (M), the nucleocapsid protein (N), and the spike protein (S) [4]. These structural proteins protect the virus from its surroundings, package the viral genome into new virions, and allow the virus to infect new cells. The second and largest class of viral proteins is the nonstructural proteins (NSPs) [4,5] with 16 members involved in viral replication, post-transcriptional modifications, and genomic processing [4,6]. The final class of proteins is the accessory proteins. This class is unique because it is unnecessary for viral replication. Instead, these accessory proteins provide the virus with additional tools to evade the host, and their presence has been correlated with the virus’s highly pathogenic characteristics [4].

Of the three classes of viral proteins, the nonstructural proteins are the primary candidates for targeted treatment because they play an essential role in viral replication and are well-conserved across different coronaviruses. However, limited literature on how these NSPs function and interact with the host is available.

Most published studies on SARS-CoV-2 NSPs have demonstrated their effects on the host cell immune response, specifically the downregulation of interferons (IFNs), a known pathway activated during viral infection [4]. For instance, NPS1 was shown to modulate different immunological response pathways, such as NF-κB, IRF_3_/IRF_7_, JAK/STAT, and others. Other proteins, such as NSP13, NSP14, and NSP15, showed potent IFN antagonism [7], whereas NSP5 stabilizes the mitochondrial antiviral-signaling protein (MAVS), resulting in an elevation of NF-κB, triggering a cytokine storm [8]. It was also shown that NSP6 modulates the interferon-stimulated response element (ISRE) promoter [9].

As these NSPs play a fundamental role in facilitating the viral life cycle, learning more about how they function can help us understand the virus’s pathogenicity.

NSP15 is a highly conserved Mn^2+^-dependent nidoviral RNA uridylate-specific endoribonuclease (NendoU) [10]. This enzyme contains three domains: an N-terminal oligomerization domain, a central domain, and a C-terminal catalytic domain. The C-terminal catalytic domain cleaves the 3’ end of uridylates to produce 2’-3’ phosphodiester and 5’-hydroxyl termini. These uridylates are generally found as poly-U repeats within the complementary single-stranded RNA template produced during viral replication. The cleavage of these poly (U) repeats is necessary because it prevents activation of pattern recognition receptors such as the retinoic acid-inducible gene I (RIG-I-like receptor) [11] and the melanoma differentiation-associated protein 5 (MDA5), a cytosolic viral RNA sensor, thereby triggering an innate immune response [7]. NSP15 also suppresses interferon alpha (IFN-α) and interferon beta (IFN-β) associated pathways to prevent the formation of cytoplasmic stress granules [16].

In the current study, we explored whether NSP15 could bind host proteins with alternative functions that benefit the virus during infection. For this purpose, we explored potential host ligands that interact with NSP15. Using mass spectrometry, protein-protein interaction techniques, and fluorescence, we discovered that NSP15 interacts with the host keratin type II cytoskeletal 1 (keratin 1), leading to changes in cytoskeletal integrity. To the best of our knowledge, this is the first study reporting a direct interaction between an SARS-CoV-2 NSP and a host protein.

## Materials and Methods

### NSP15 cloning and expression

The gene *NSP15* (IDT, Coralville, IA, USA) was cloned into pET-28 using *Bam*HI and *Hind*III (NEB, Ipswich, MA, USA) restriction enzymes. The recombinant NSP15 protein was produced by transforming the construct into *E. coli* BL-21. NSP15 was purified by affinity chromatography using a Ni-NTA resin and following the manufacturer’s instruction (Qiagen, Hilden, Germany).

### Protein-protein interaction

A lysate of Vero E6 cells (ATCC-CRL 1587) was prepared by sonicating the cells in a buffer solution containing Tris-HCl (50 mM, pH 7.5), DTT (1 mM), NP-40 (0.05%), and NaCl (100 mM). After clearing the sample by centrifugation, the lysate was mixed with 100 ∝g of recombinant NSP15 and rotated overnight at 4 °C. The mixture was then loaded on a Ni-NTA resin, and the NSP15 was eluted according to the manufacturer’s instructions. The elution fraction was loaded onto an SDS-PAGE gel, and after resolution, the gel was silver-stained for mass spectrometry analysis. The bands of interest were excised and stored in 20% acetonitrile for further processing.

### Proteomic analysis

In-gel trypsin digestion was used to generate peptides for LC-MS analysis. Bands were cut into 1 mm pieces, washed twice with 100 µL of 50 mM ammonium bicarbonate for 10 min each, then dried by treating with 100 µL of acetonitrile for 10 min, and finally dried in a vacuum centrifuge (Centrivap, Labconco, Kansas City, MO, USA). Reduction and alkylation were carried out with 50 µL of 10 mM DTT for 35 min at 65 °C, followed by removal of the excess, and then 30 µL of 55 mM iodoacetamide for 30 min at RT in the dark. The supernatant was removed again, and the gel pieces were washed twice with 50 mM ammonium bicarbonate for 10 min, then dried as above with acetonitrile and Centrivap before being dissolved in 10 µL of trypsin (10 ng/µL, NEB). The trypsin solution was absorbed into the dried gel pieces, then 50 mM ammonium carbonate was added to cover the gel pieces, and samples were incubated overnight at 37°C. The supernatant was transferred to microcentrifuge tubes, and gel pieces were extracted twice for 15 min with 50 µL of 50% acetonitrile and 5% formic acid. Extracts were pooled with the initial supernatant, dried, dissolved in 15 µL 0.1% trifluoroacetic acid, and transferred to the LC sample plate.

Analysis was conducted on an Orbitrap Fusion Lumos MS platform coupled to an Easy-nLC 1200 system (ThermoFisher, Waltham, MA, USA) using an in-house 100 µm ID x 100 cm column (Dr. Maisch 1.9 µ C18). The column was equilibrated at 600 bars for a total volume of 6 μL. A gradient of mobile phase A (water and 0.1% formic acid) and B (80% acetonitrile with 0.1% formic acid) at 0.4 µL/min, 2-27% B from 1-35 min followed by 27-45% B over 10 min, 45-95% B over 4 min, and 10 min at 95% was used (1 h total). Data acquisition was performed using a data-dependent method in the ion trap, with MS2, using a positive ion spray voltage of 2100, a transfer tube temperature of 325**°**C, and a default charge state of 2. Survey scans (MS1) were acquired at a resolution of 120 K across a mass range of 375 – 1500 *m/z*, with RF 30, an AGC target of 4e5, and a maximum injection time of 50 ms in profile mode. For MS2 scans, there was an intensity threshold of 5e3, charge state 2–5, dynamic exclusion 20 seconds with 10 ppm tolerances with a 1.2 *m/z* window and using CID fragmentation with 35% collision energy, 10 msec activation, normal scan rate, auto scan range, 100% AGC target, and maximum injection time of 54 ms in centroid mode.

All data files were processed with Protein Discoverer 2.5. Spectrum files were recalibrated, and features were extracted with Minora. Searches were carried out using Sequest HT with Uniprot *Cercopithecus sabaeus* UP000029965, precursor mass tolerance of 20 ppm and fragment mass tolerance of 0.5 Da, static carbamidomethyl modification, and K, M, P oxidation, and dynamic S, T, Y phosphorylation modifications. Decoy database strict and relaxed FDR targets were 0.01 and 0.05 based on q value, and the data were exported to Excel for further analysis.

### Immunoblotting

An SDS-PAGE was prepared, and proteins were separated and transferred onto a nitrocellulose membrane (BioRad, Hercules, CA, USA) at 300 mA for 30 min. Then, the membrane was transferred to a glass chamber and blocked with 3% BSA in PBS overnight at 4°C. The next day, BSA was discarded, and a mouse anti-cytokeratin 1 primary antibody (1:1000, Novus Biologicals) was added to the chamber and rotated for 3 h at RT. The primary antibody was removed, and the membrane was washed with PBS + 0.05% Tween-20 (×3) for 10 min each. Then, 1:5000 Alexa Fluor 680 goat anti-mouse secondary antibody (ThermoFisher) was added and rotated for 1 h at RT in darkness. The secondary antibody was removed, and the membrane was rewashed × 3 with PBS + 0.05% Tween-20 for 10 min each. The solution was discarded, and the membrane was scanned using an Odyssey^®^ imaging system (Lycor Bio, Lincoln, NE, USA) to visualize the bands at 680 nm.

### Protein purification and incubation

Vero E6 cells were pelleted, incubated with lysate buffer, and sonicated at 9 V for 30 sec until the pellet was dispersed. The tube was then centrifuged at 2000 × *g* for 10 min, and the supernatant containing the intracellular proteins was supplemented with purified his-tagged NSP15 protein (100 μg) and left to rotate overnight at 4°C.

### Immunoprecipitation

The Vero E6 cell pellet was incubated with lysate buffer and sonicated at 9 V for 30 sec until the pellet was dispersed. The tube was then centrifuged at 3,000 × *g* for 5 min to pellet the debris, and the supernatant was transferred to a new tube. A reaction containing 200 µg of NSP15, Vero E6 cell lysate, and 1 mM PMSF was incubated at 4 °C for 1 h in a tube. Mouse anti-his-tag antibody (1:5000, Qiagen) was then added to the tube and rotated for 1 h at 4°C. The protein mixture was then passed through a protein G column (GE, Boston, MA, USA), washed with PBS, and eluted using 0.2 M glycine-HCl, pH 2.2. The concentration of the proteins was measured with a Bradford assay using a spectrophotometer at 595 nm.

### Immunofluorescence

Vero E6 cells (1×10^5^) were plated on sterilized glass coverslips in a 24-well plate using MEM media (Invitrogen, Waltham, MA, USA) supplemented with 10% fetal bovine serum (FBS) (Invitrogen), sodium pyruvate (Invitrogen), non-essential amino acids (Invitrogen), penicillin/streptomycin (Invitrogen), and amphotericin B (Invitrogen). The cells were incubated at 37 °C supplemented with 5% CO_2_ overnight. The next day, the culturing media were discarded, and cells were incubated with PBS for 15 min at 37°C twice. The cells were then fixed with 4% paraformaldehyde (PFA) for 10 min at 37°C and rinsed with PBS twice. Cells were then incubated with PBS (×3) for 15 min at 37°C. Goat serum supplemented with saponin was used to permeabilize cells for 30 min at RT. Anti-cytokeratin 1 primary antibody (1:1000) was added for 30 min at RT. Coverslips were then rinsed (×3) with PBS and incubated in PBS for 10 min at RT. Cells were then permeabilized again using saponin for 30 min at RT. Goat FITC-labeled secondary antibody (1:500, ThermoFisher) was added for 30 min at RT in darkness. The coverslip was rinsed with PBS (×3) and incubated in PBS for 10 min at RT. Coverslips were then rinsed with ddH_2_O, mounted onto a glass slide with DAPI-fluorosafe, and stored in the dark overnight at RT. Samples were sealed with nail polish, and the images of the cytoskeleton and nuclei were then visualized and captured using fluorescence microscopy.

FITC was excited using a 488 nm laser to detect keratin 1, and the sample was visualized using a Zeiss LSM780 confocal microscope equipped with 40X and 60X oil-immersion objective lenses. Images were acquired using ZEN imaging software.

### Protein profection into Vero E6 cells

Vero E6 cells (1×10^5^ cells) were dispensed onto sterilized circular glass coverslips in a 24-well plate, following protocols previously published in our lab [12]. The next day, 6 μL of Profect-P1 (Targeting Systems, El Cajon, CA, USA), 1.8 μg recombinant NSP15, and 150 μL MEM were combined in a single tube and incubated for 20 min at RT, then 600 μL MEM was added to the tube. Cells were rinsed twice with MEM, and 200 μL of the Profect-P1 mixture was added to each well and incubated at 37°C. Time points at 1, 2, and 3 h were collected. The media was aspirated, rinsed with PBS, and fixed with 4% paraformaldehyde for 30 min.

### MTT cytotoxicity assay

THP-1 cells (1×10^5^ cells/well) were dispensed into a 96-well plate, and 100 μL of complete media was added to each well and incubated at 37°C, supplemented with 5% CO_2_ overnight and following published protocols.

Culturing media was aspirated from each well, and 100 μL of Profect solution control, Profect solution without NSP15 or 10% SDS, was added to the wells, and the plate was incubated at 37°C. At the end of each measured time point (20, 40, 60, 120, 180, 360, and 540 min), 10 μL of a sterile MTT solution (5 mg dissolved in PBS, Sigma) was added to the wells. After 4 hours, 100 μL of the dissolving buffer was added to each well to terminate the reaction. Experiments were performed in triplicate.

The following day, the plate was read at 570 nm using a spectrophotometer (Epoch, Agilent).

### Phalloidin-FITC actin staining

Phalloidin-FITC (13 μL from a stock of 50 mg/mL) was added for 40 min to a coverslip containing fixed cells. The coverslip was then rinsed with PBS and incubated with PBS ×3 for 10 min each. The coverslips were then mounted onto a glass slide with 3 μL DAPI-fluorosafe and left overnight at RT. The next day, samples were sealed with nail polish. The sample was then visualized using a Zeiss LSM780 confocal microscope with 40X and 60X oil-immersion objective lenses. Images were acquired using ZEN imaging software.

## Results

### Identification of keratin as a ligand of NSP15

To identify potential NSP15 substrates in the host cell, recombinant NSP15 was incubated with Vero E6 cell lysates and then purified using a Ni-NTA resin to pull down potential ligands.

The silver staining showed two bands at 45 kDa and 60 kDa (black arrows, Fig 1, lane 4), which were absent in the positive control containing only NSP15 (orange arrow, Fig 1, lane 3). This observation indicated a possible interaction with at least two host proteins. The protein bands were excised and identified by liquid chromatography-mass spectrometry (LC-MS).

**Fig 1.**
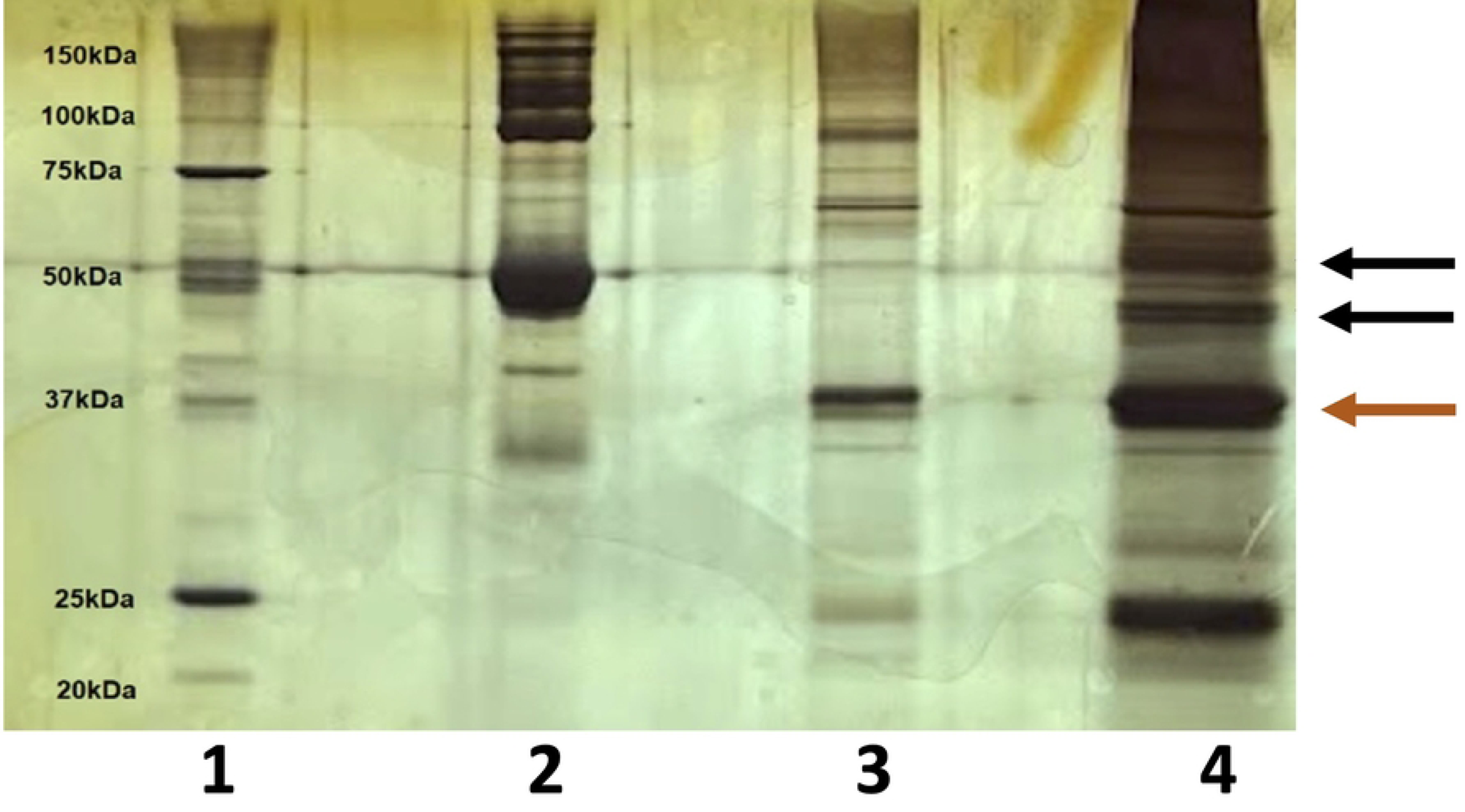
Protein-protein interaction between NSP15 and host proteins. Recombinant NSP15 was exposed to a lysate of Vero E6 cells overnight at 4°C. The mixture was purified using Ni-NTA resin, and the eluate was resolved on a 10% SDS-PAGE gel and stained with silver. Arrows point to the new protein bands detected by LC-MS. Lanes, 1, Protein marker; 2, 3% BSA (negative control); 3, Recombinant NSP15; 4, Vero E6 cell lysate incubated with recombinant NSP15. Two bands (black arrows) around 45 and 60 kDa in lane 4 indicate potential Vero E6 cell proteins interacting with NSP15, whereas the orange arrow indicates the migration of NSP15.

### Identification of the protein interacting with NSP15

The LC-MS results revealed that the two unknown bands in the gel were keratin (Table 1). Furthermore, the various types of keratins appearing as hits also suggested that NSP15 could likely bind to a conserved region of the keratin family.

**Table 1.** Liquid chromatography-mass spectrometry results show the protein identities corresponding to the two unknown bands.

| Access # | Protein (*Homo sapiens*) | MW (kDa) | Peptide # | Coverage (%) | Abundance |
| --- | --- | --- | --- | --- | --- |
| P04264 | Keratin, type II cytoskeletal 1 | 66 | 26 | 52 | 2.55E+08 |
| P35527 | Keratin, type I cytoskeletal 9 | 62 | 19 | 53 | 2.05E+08 |
| P13645 | Keratin, type I cytoskeletal 10 | 58.8 | 20 | 49 | 6.43E+07 |
| P35908 | Keratin, type II cytoskeletal 2 epidermal | 65.4 | 23 | 49 | 5.96E+07 |
| P00367 | Glutamate dehydrogenase 1, | 61.4 | 13 | 29 | 4.56E+07 |
| Q7Z794 | Keratin, type II cytoskeletal 1b | 61.9 | 2 | 2 | 2.12E+07 |
| P13647 | Keratin, type II cytoskeletal 5 | 62.3 | 7 | 13 | 3.66E+06 |
| P48668 | Keratin, type II cytoskeletal 6C | 60 | 9 | 19 | 2.53E+06 |

Mass spectrometry results suggest that various isoforms of cytoskeletal keratin between the sizes of 58.8 to 66 kDa interacted with NSP15 (Table 1).

### Validation of keratin expression in Vero E6 cells

Before validating the interaction between NSP15 and keratin, we demonstrated that the keratin type II cytoskeletal 1 identified by LC-MS was expressed in the cell line from which we isolated our sample. This was accomplished by mixing the Vero E6 cell lysate and incubating the lysate with NSP15. An immunoblot was performed using a sample obtained from the Ni-NTA resin and resolved in an SDS-PAGE. The gel was transferred to a nitrocellulose membrane and probed with a mouse anti-cytokeratin 1 primary antibody (Fig 2). The immunoblot results revealed an approximate 60 kDa band consistent with the size of the keratin. This confirmed that keratin identified by LC-MS was indeed expressed in Vero E6 cells.

**Fig 2.**
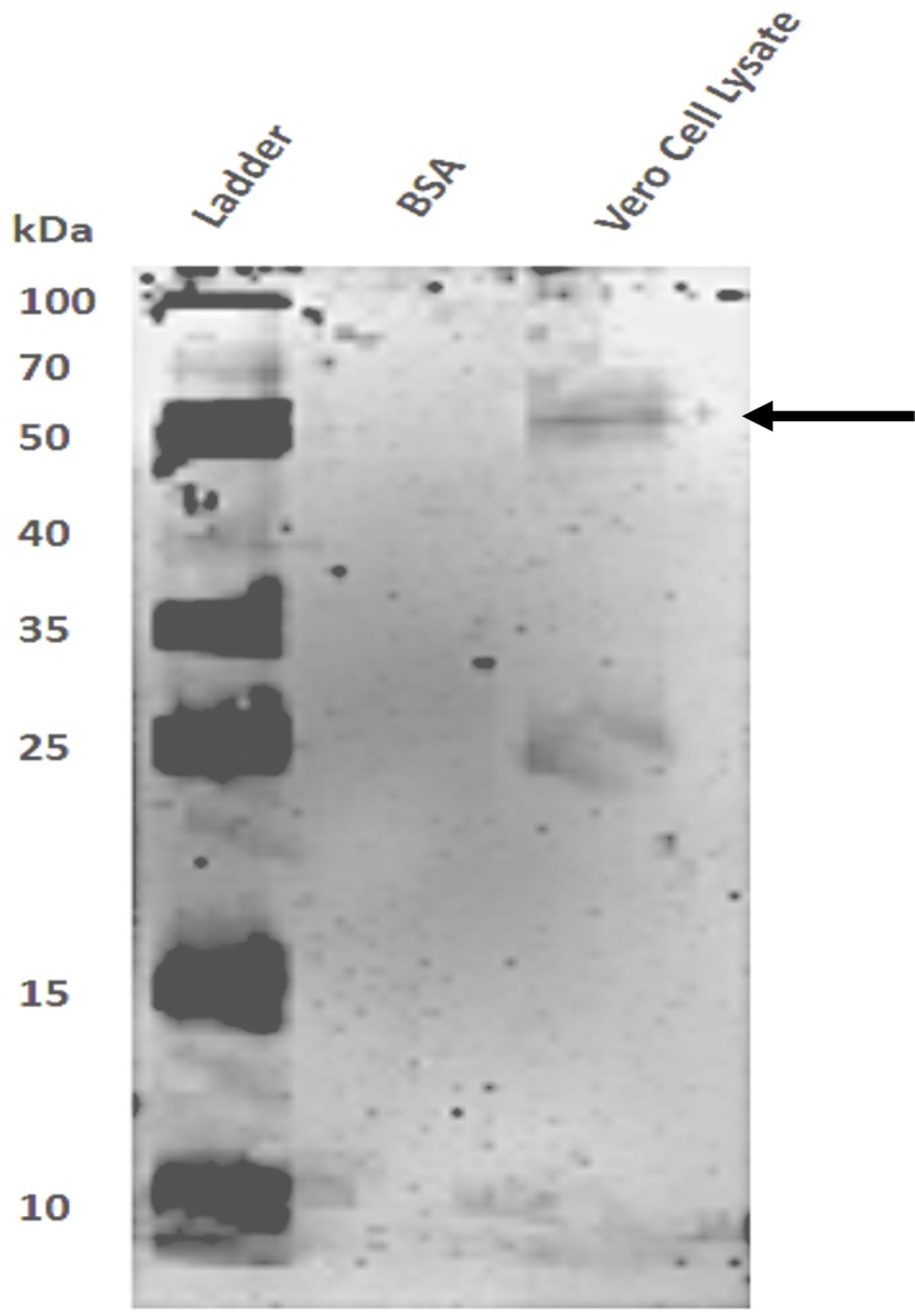
Confirmation of keratin expression in Vero E6 cells. Expression of keratin in Vero E6 cells was validated by immunoblot. Lanes: 1, Protein ladder; 2, BSA (negative control); 3, Vero E6 cell lysate. The membrane was exposed to a mouse anti-cytokeratin primary antibody and a goat anti-mouse fluorophore-conjugated secondary antibody (680 nm). Results showed a band around 60 kDa consistent with the size of keratin. Another unknown 25 kDa protein was also detected.

### Validation of NSP15-keratin 1 interaction by immunoprecipitation and immunoblotting

The interaction between NSP-15 and keratin 1 was validated using immunoprecipitation. Once again, a cell lysate was prepared from Vero E6 cells and incubated with NSP15. Then, a mouse anti-His-tag antibody was added to the reaction and loaded onto a Protein G column to bind the IgG anti-His-tag antibody selectively. The eluate was collected, and proteins were separated by electrophoresis and transferred onto a nitrocellulose membrane. An immunoblot was performed using the same anti-cytokeratin primary antibody to validate the keratin expression (Fig 3). These results validated NSP15 binding to keratin.

**Fig 3.**
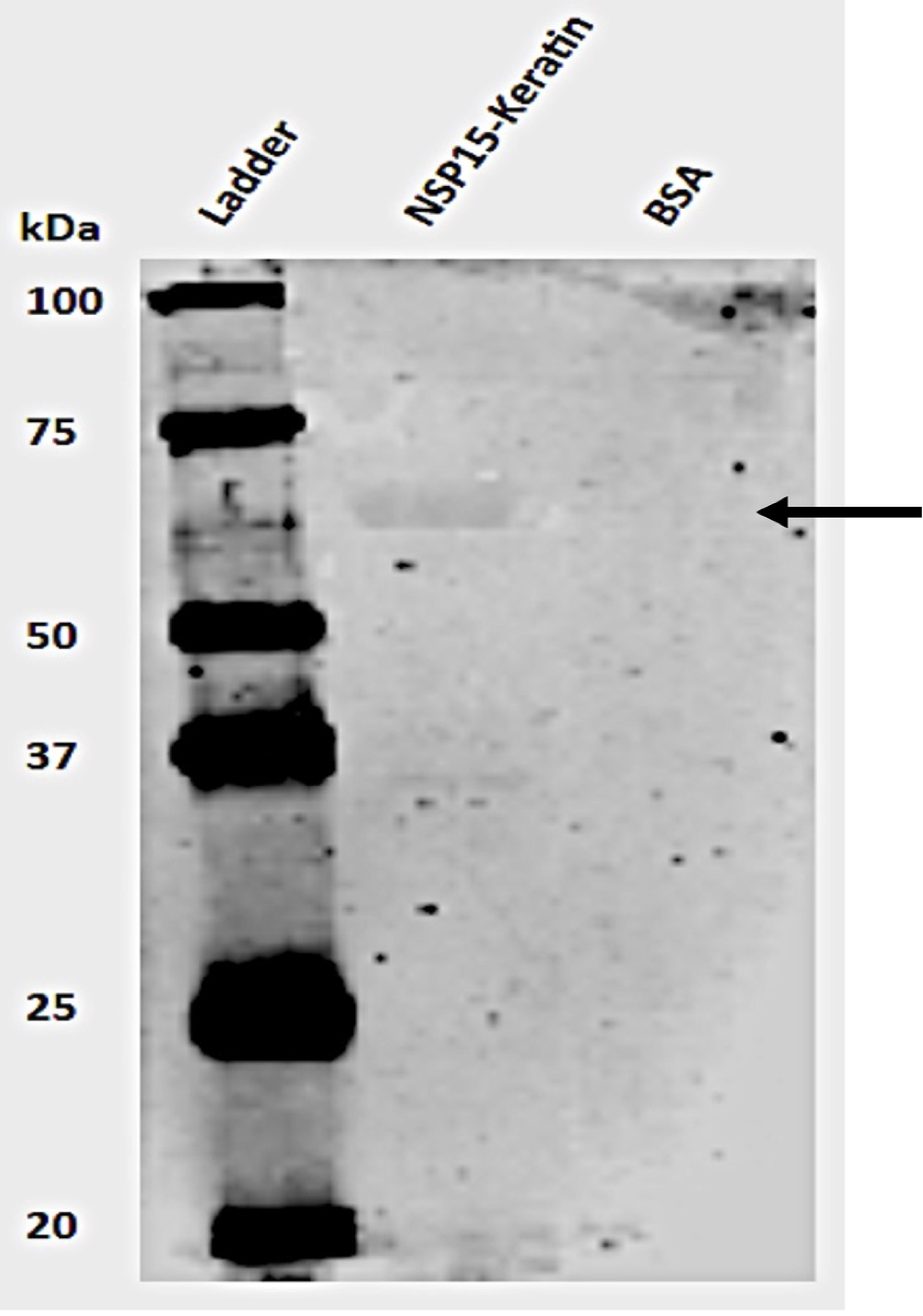
Validation of protein-protein interaction between keratin and NSP15. An immunoblot showed the interaction of recombinant NSP15 with keratin. NSP15 was pulled out with mouse anti-His-tag antibodies bound to Protein G, and the membrane was probed with an anti-cytokeratin antibody. Lanes, 1, Protein ladder; 2, NSP15-Vero E6 lysate-antibody mixture; 3, 3% BSA (negative control). The black arrow points to the keratin.

Further controls were performed to rule out that the recombinant NSP15 or keratin alone could not bind the Protein G. An immunoblot showed no bands when only NSP15 (Fig 4A) or Vero E6 cell (Fig 4B) lysates were passed through the Protein G.

**Fig 4.**
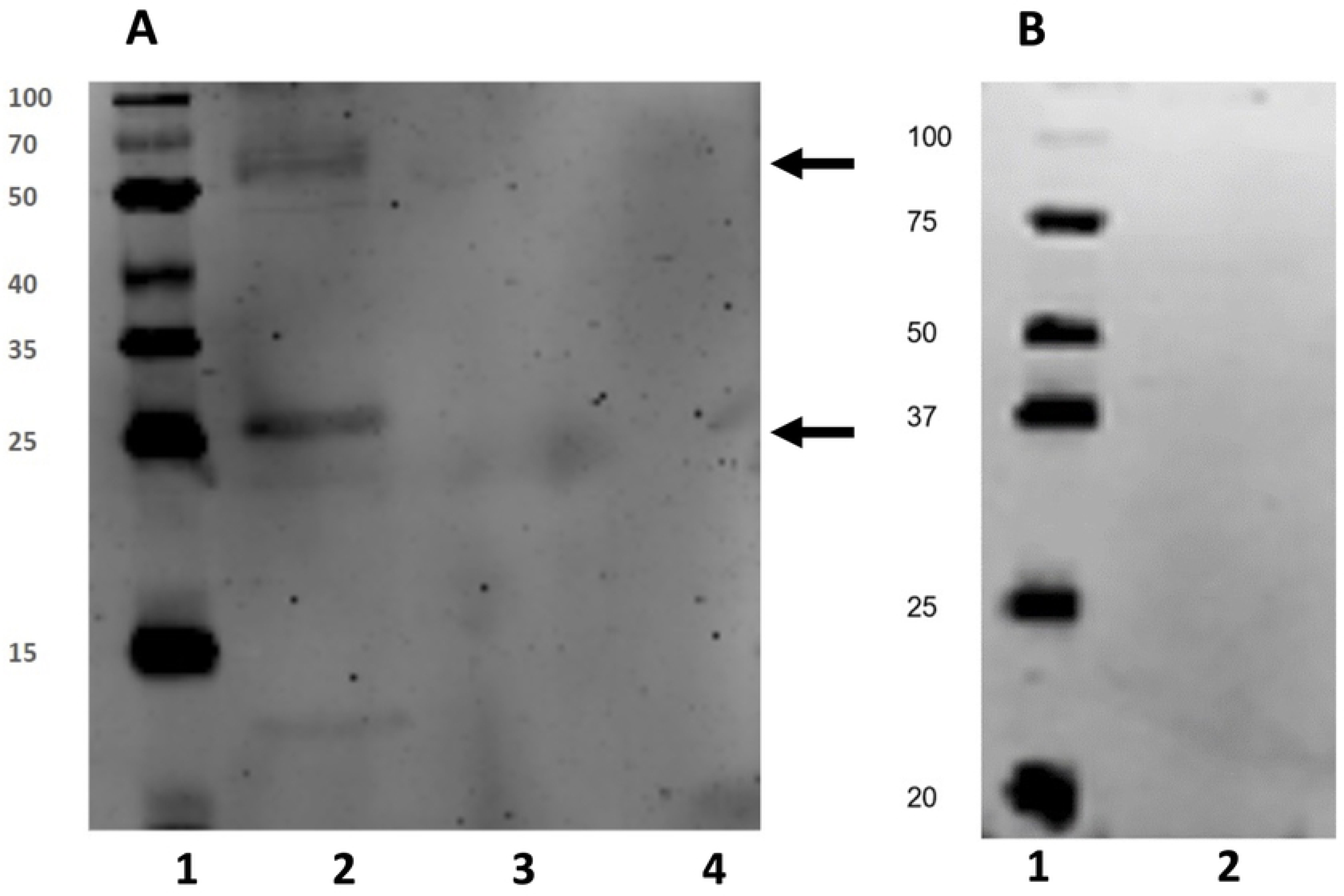
NSP15 and the Vero E6 cell lysate did not bind Protein G. (A) No bands were identified when the recombinant NSP15 was tested. Lanes: 1, Protein ladder; 2, NSP15-keratin eluate showed a band at 60 kDa, consistent with the size of keratin (upper arrow) and an unknown 25 kDa protein (lower arrow); 3, NSP15 eluate; 4, 3% BSA (negative control). (B) No bands were observed when the cell lysate was tested. Lanes: 1, Protein ladder; 2, Vero cell lysate.

### Visualization of keratin intermediate filaments in Vero E6 cells

Having shown that NSP15 binds to the cytoskeletal keratin, the next goal was to identify a potential function for this protein-protein interaction. Thus, we stained the keratin cytoskeletal network with the same mouse anti-cytokeratin primary antibody and goat anti-mouse FITC-conjugated secondary antibody. This staining revealed the intricate network of the keratin intermediate filaments within the Vero E6 cell (Fig 5).

**Fig 5.**
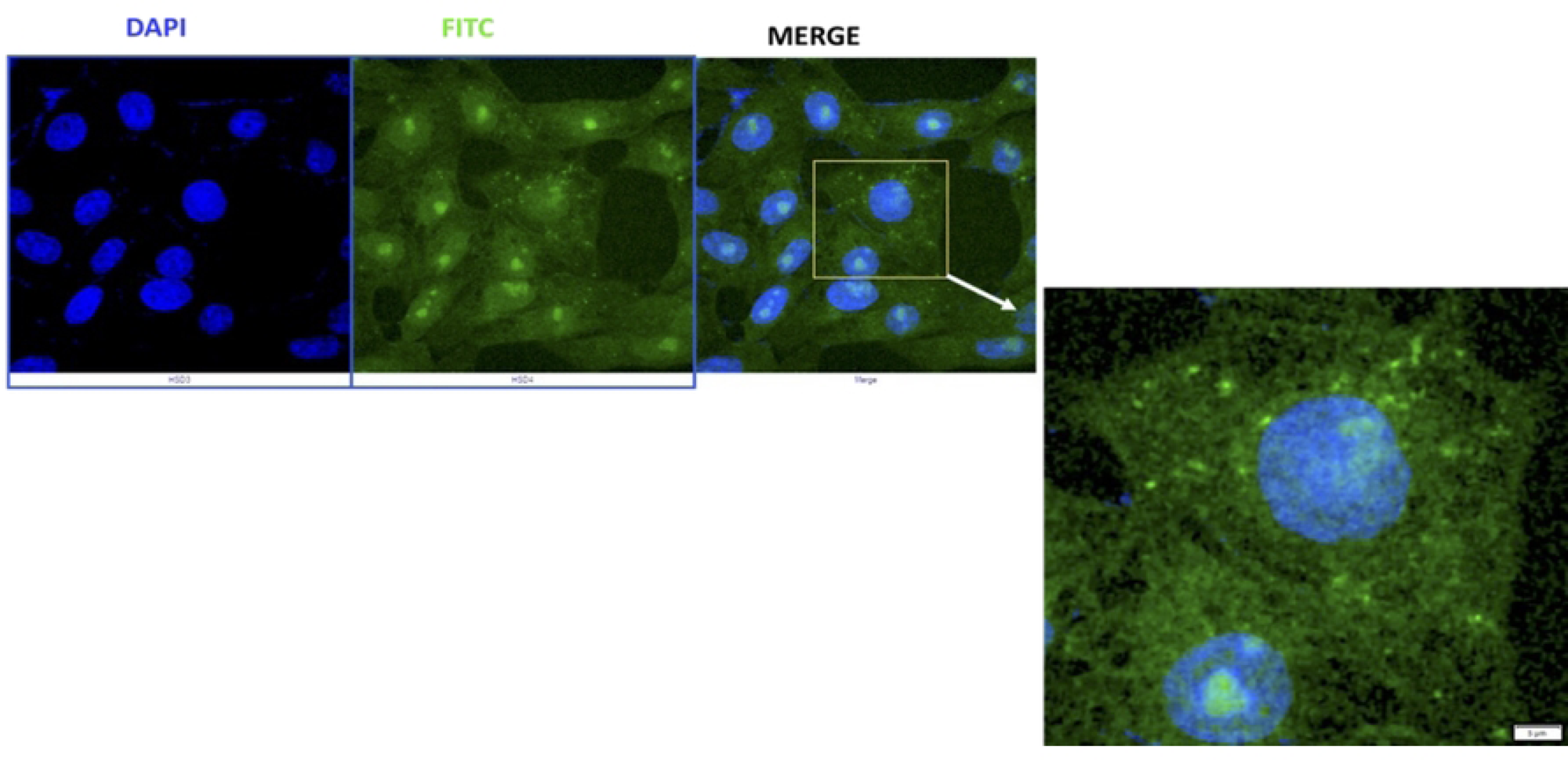
FITC-labeled cytokeratin network in Vero E6 cells. The cytokeratin network in Vero E6 cells was labeled using a mouse anti-cytokeratin primary antibody and a goat anti-mouse FITC secondary antibody (green). A DAPI stain was used to visualize the nuclei (blue). A fibrous cytokeratin network spanning the cell is visible in Vero E6 cells. Scale bar = 5 mm.

### Impact of NSP15 and keratin on cytoskeletal integrity in Vero E6 cells

To evaluate the impact of the NSP15-keratin interaction, recombinant NSP15 was internalized into the Vero E6 cells via profection. A time-dependent collapse of the cytoskeletal network was observed when these cells were analyzed under the fluorescent microscope (Fig 6). The longer the cells were incubated with the viral protein, the more significant the disruption in cytoskeletal structural integrity was observed (Fig 6A). This suggests that the interaction between NSP15 and keratin collapses keratin intermediate filaments in infected cells. As a result of this collapse, an increase in cytotoxicity was measured by an MTT assay (section 3.7, Fig 7).

**Fig 6.**
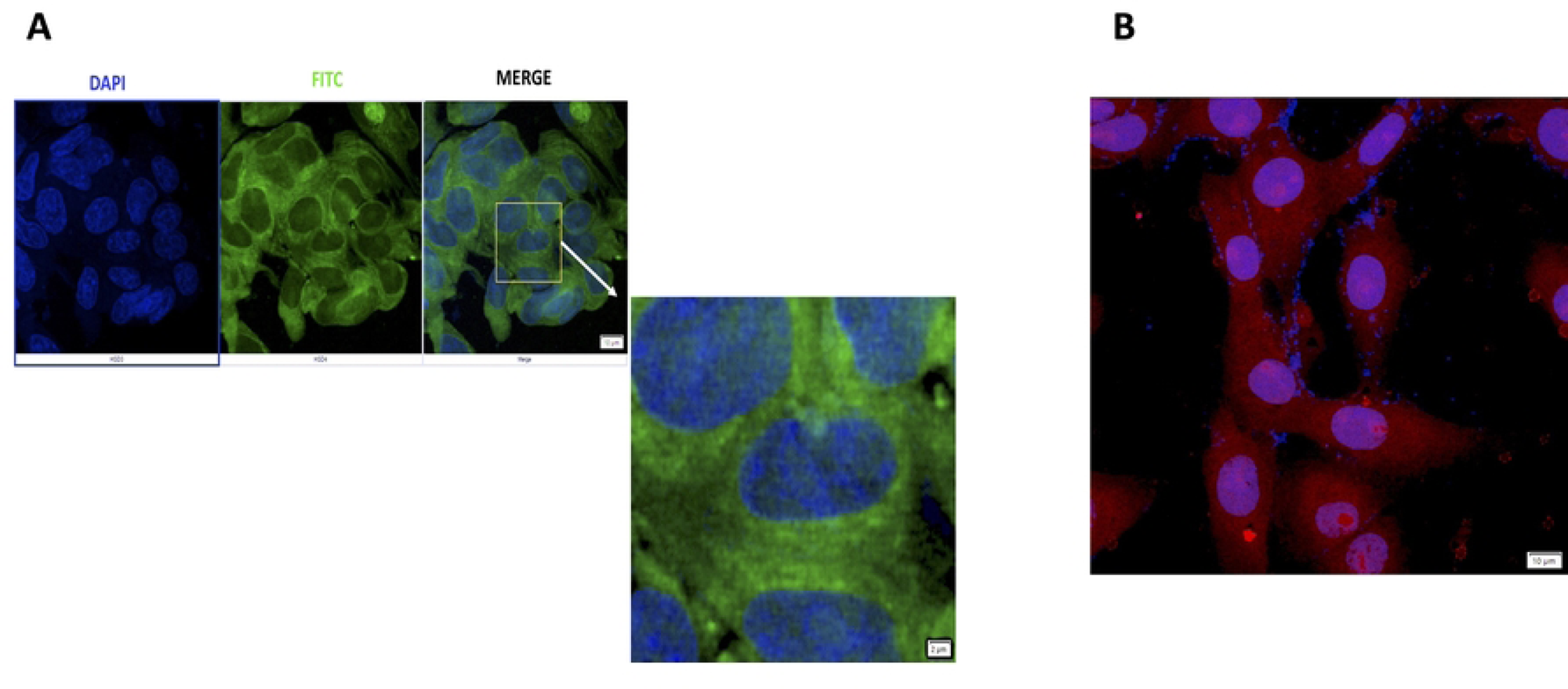
FITC-labeled cytokeratin network in Vero E6 cells profected with recombinant NSP15. (A) Cytokeratin network in Vero E6 cells after 3 h post-incubation with NSP15 was labeled using mouse anti-cytokeratin primary antibody and goat anti-mouse FITC secondary antibody (green). A DAPI stain (blue) was used to visualize the nuclei. In the merged image, the fibrous patterns of the cytokeratin network are lost, and the green fluorescent signal diffuses throughout the cell. Scale bar = 2 μm. (B) A control experiment for the profection protocol was used to confirm a successful internalization of recombinant red fluorescent protein into Vero E6 cells. Scale bar = 10 μm.

**Fig 7.**
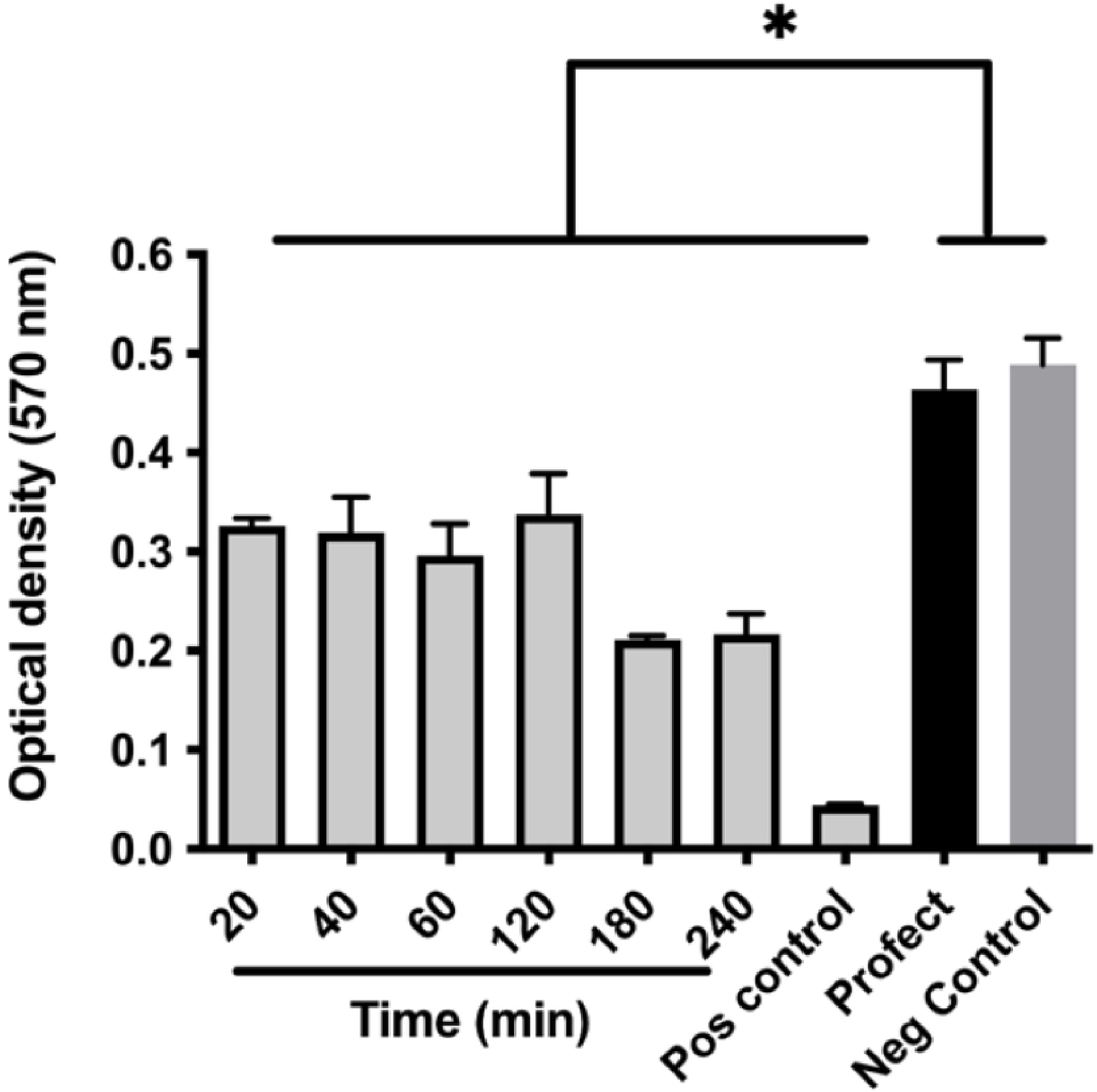
Cytotoxic effects of NSP15. MTT cytotoxicity assay was implemented to measure the cell viability over time in Vero E6 cells profected with NSP15. Untreated and Profect (no NSP15) were used as negative controls, whereas SDS (10%) was used as a positive control. Error bars indicate the mean of three independent experiments ± standard deviation.

### Cytotoxicity of the profected NSP15

As mentioned above, a kinetic study was performed to evaluate the cytotoxicity of the profected NSP15. Results showed a time-dependent reduction in Vero E6 cell viability when NSP15 was profected into the Vero E6 cells (Fig 7). After 20 minutes of incubation with NSP15, there was already a statistically significant reduction in cell viability (ANOVA, p<0.01). This indicated that the time scale for disrupting the keratin network was rapid and potent. Although the physiological concentrations of NSP15 are likely lower than the ones utilized in the current study, it is reasonable to suggest that as the virus replicates over time within the infected cell and the viral proteins begin accumulating, this could lead to the disassembly of the keratin intermediate filaments and help to release the viral progeny from the host cell.

### NSP15 does not interact with the actin cytoskeleton

Another structural component of the cytoskeleton is the actin filaments. Thus, to determine whether or not NSP15 also affects the actin cytoskeleton, a set of immunofluorescence experiments was performed after exposure of Vero E6 cells to NSP15, but labeling the actin cytoskeleton instead of the keratin. When NSP15 was introduced into Vero E6 cells, no structural changes in the actin cytoskeleton were observed, even after 3 hours of incubation (Fig 8). In summary, unlike the keratin network, the actin filaments are completely unaffected by the presence of NSP15. This suggests that NSP15 does not disrupt the actin cytoskeleton.

**Fig 8.**
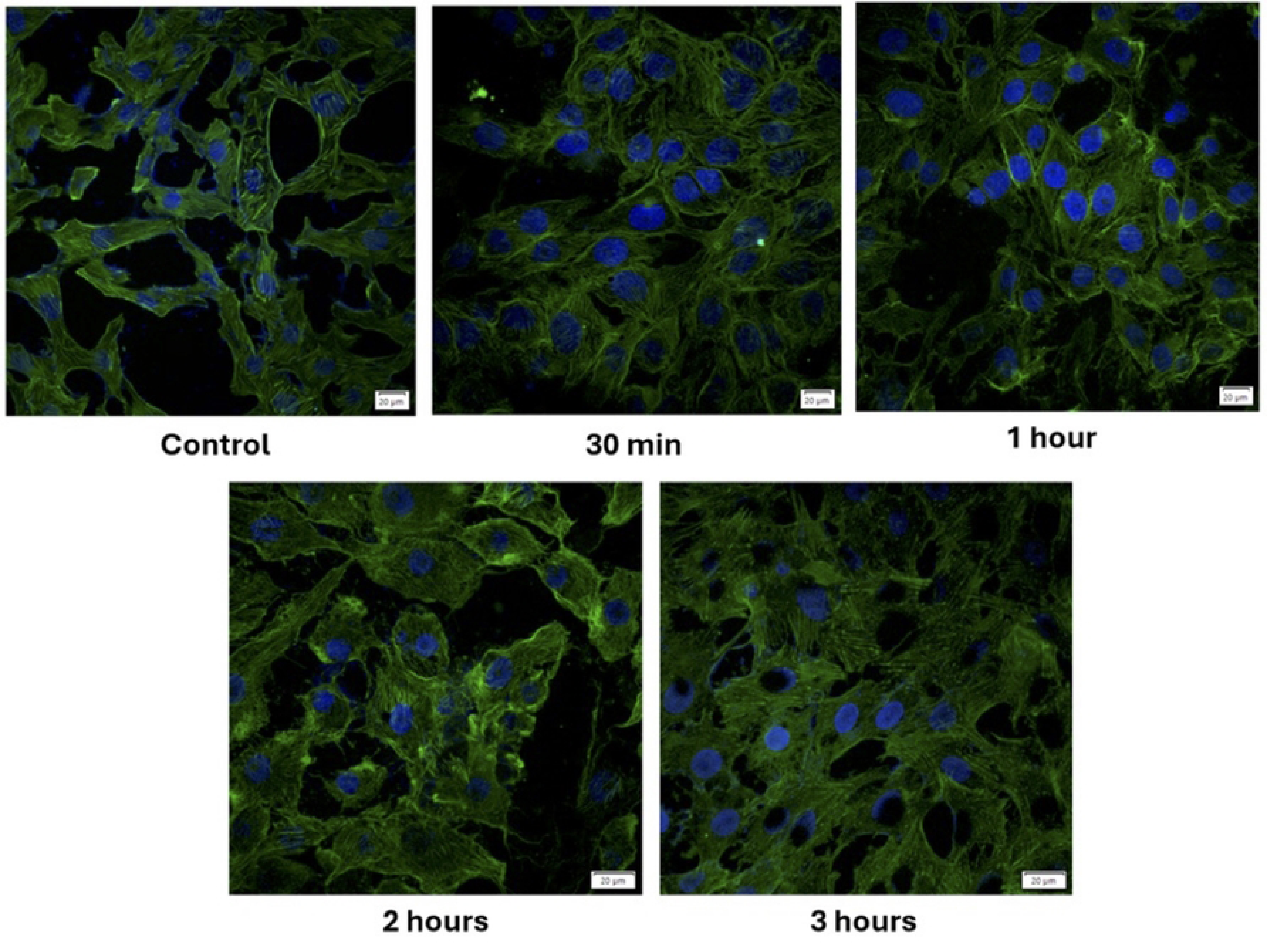
Fluorescence microscopy reveals that NSP15 does not affect the actin cytoskeleton. Fluorescent images showing the actin cytoskeleton labeled with phalloidin-FITC (green). Vero E6 cells transfected with NSP15 for up to 3 hours show no structural changes to the actin cytoskeletal network or the shape of the nuclei labeled with DAPI (blue). Scale bar = 20 μm.

### Co-localization of NSP15 and keratin in Vero E6 cells

Having demonstrated that the transfection of NSP15 into Vero E6 cells leads to a disruption of the keratin cytoskeleton, we performed a co-localization experiment for NSP15 and the cytokeratin. After transfection of NSP15 into Vero E6 cells for 3 hours, NSP15 (Fig 9C, red) and keratin (Fig 9D, green) co-localized as shown by the presence of yellow spots produced by the overlapping signals of NSP15 (red) and keratin (green) (Fig 9).

**Fig 9.**
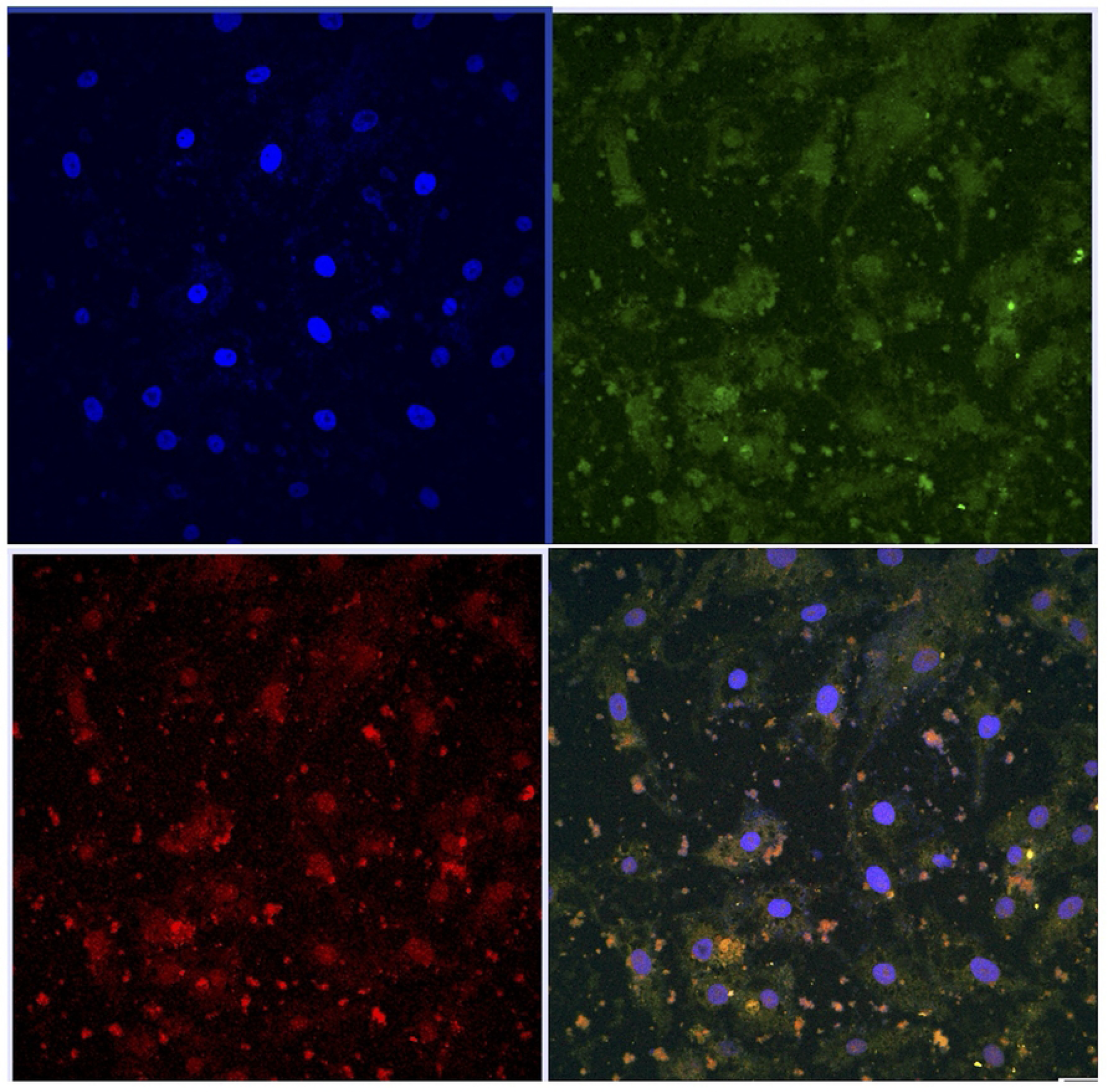
Co-localization of NSP15 and keratin in Vero E6 cells. A co-localization experiment validated that NSP15 (red) interacted with keratin (green) within the cell by overlaying the two fluorescent images (yellow). Scale bar = 20 μm.

In summary, the results indicated that this novel protein-protein interaction between NSP15 and keratin in Vero E6 cells disrupts the structural integrity of intermediate filaments, directly reducing cell viability.

A quantification of the co-localization using the fluorescence microscope was performed as shown in Fig 10, which showed that both proteins follow the same pattern of intensities.

**Fig 10.**
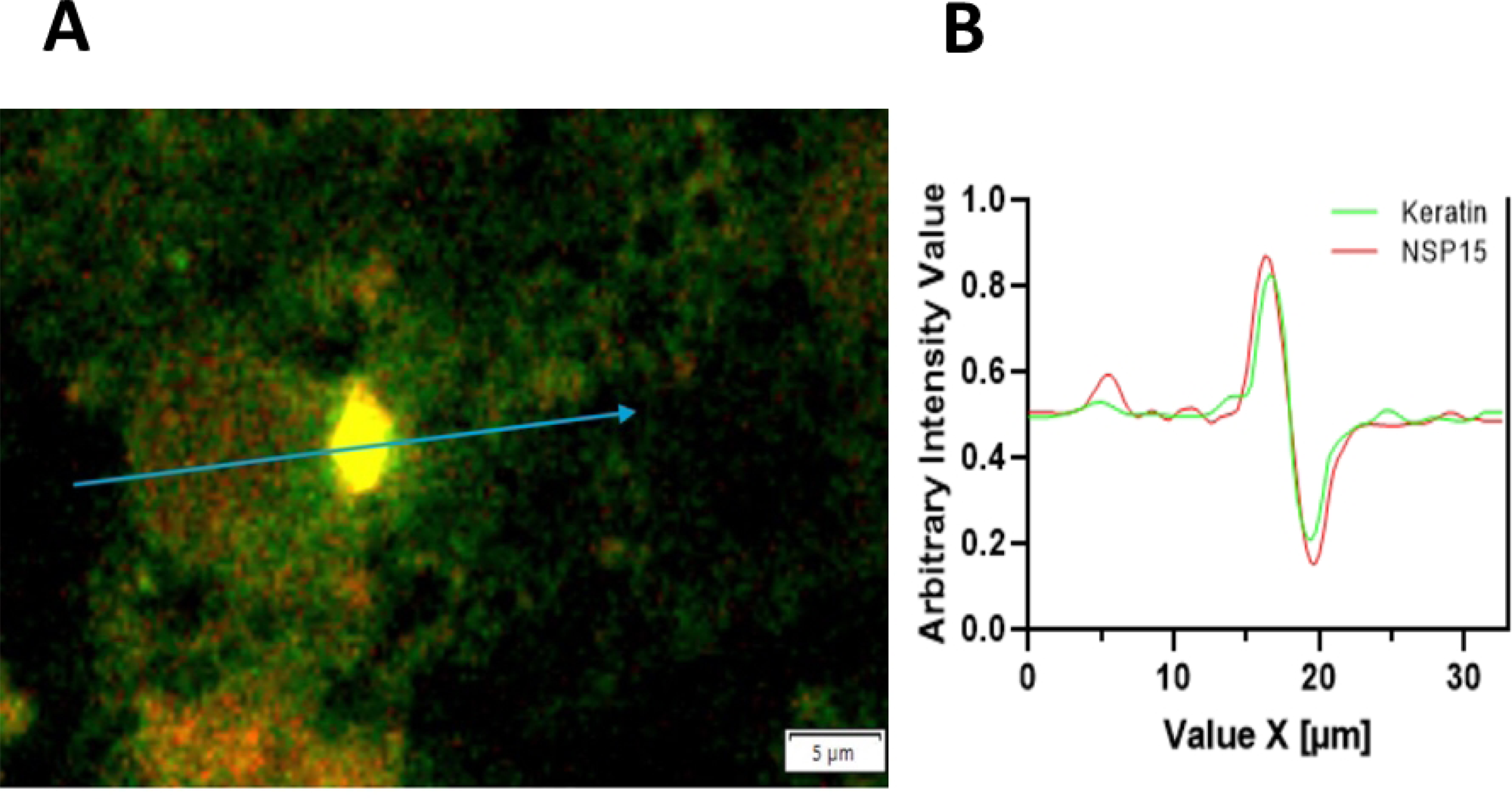
Co-localization quantification of keratin (green) with NSP15 (red) foci. (A) The intensity was quantified along the blue arrow, and the results are shown in the green and red channels in (B). DAPI (blue) was excluded for quantification. Scale bar = 5 μm.

## Discussion

The structural composition of the SARS-CoV-2 virus and the mechanisms it employs to infect its host are well known. Still, much remains to be understood about the intricacies of viral pathogenesis and how viral proteins interact with the host to promote viral replication. The nonstructural proteins are essential to the survival and propagation of the virus. The loss of these nonstructural proteins significantly reduces viral pathogenicity as they are required for viral replication [13]. As the functions of the NSPs appear to be well conserved across different coronaviruses, we have a basic understanding of how they contribute to viral replication and promote the viral life cycle. However, assuming that these viral proteins serve only a single function would be shortsighted, as proteins can participate in multiple pathways and have unintended off-target effects. Due to the limited size of the viral genome, it is not uncommon for the same viral proteins to play various roles in the viral life cycle, some of which may be unknown. By acknowledging this, researchers can better focus their efforts on identifying the protein-protein interactions in which SARS-CoV-2 is involved and the key interactions responsible for its enhanced pathogenicity.

In this study, we aimed to identify and validate the protein-protein interaction between NSP15 and its new substrate, the cytoskeletal keratin. Keratin organization is one of the protein systems that maintains structural stability and flexibility and ensures the mechanical resistance of epithelial cells. Keratin members vary in their sizes between 40–70 kDa and are divided into two main groups: the smaller or low molecular weight acidic type I (40–55 kDa) and the larger or high molecular weight neutral-basic type II (56–70 kDa) [14]. Here, we showed the results of the interaction between the neutral-basic keratin and NSP15. Also, we showed that upon binding to NSP15, the keratin cytoskeleton collapses, consistent with other studies that observed the same phenomenon when foreign proteins are introduced [15].

The currently available literature suggests that NSP15 primarily mediates host immune evasion by cleaving poly (U) repeats on the complementary RNA strand and suppressing IFN-α- and IFN-β-associated pathways. However, the literature does not identify any other specific protein-protein interactions between NSP15 and the host.

Although there are no prior recordings linking NSP15 or any other SARS-CoV-2 viral protein to keratin, there have been documented cases of viral proteins interacting with keratin in other viruses, such as the Human Papillomavirus 16 (HPV-16), Herpes Simplex Virus (HSV), and influenza H1N1. In HPV-16, the C-terminus of the viral E4 protein, which is usually involved in viral replication, interacted with keratin [16]. This interaction led to the collapse of the intermediate filaments in the cell and subsequently resulted in cell lysis [16]. Similar results were observed in our study when NSP15 was introduced into host cells, resulting in cytotoxicity. This is probably related to the collapse of the keratin cytoskeleton. In the case of HSV, a viral US3 protein kinase could phosphorylate keratin-17 [17]. This keratin 17 phosphorylation reduced the filamentous keratin’s staining, suggesting that the addition of phosphate to keratin disrupted the intermediate filament structure. This disruption was also linked to cytopathic effects such as cell rounding [17]. Finally, in H1N1 infections, elevated phosphorylation of keratin-8 in alveolar epithelial A549 cells was associated with changes in cell morphology [18]. The observed results were similar to the HSV findings, where increased phosphorylation was positively correlated with disassembly and elevated viral replicative efficiency. These three cases demonstrate that viral proteins interact with keratin and that all these interactions disrupt the keratin intermediate filament cytoskeleton.

Natural products have also been shown to have potential inhibitory effects against NSP15 [19]. For example, epigallocatechin gallate (EGCG), a natural compound derived from green tea extracts, inhibited NSP15 by preventing the cleavage of uridylates from the growing RNA strand. Interestingly, EGCG completely inhibited SARS-CoV-2 replication at concentrations as low as 1 mg/mL [20]. Another compound is tipiracil, an uracil derivative that inhibits thymidine phosphorylases, and it is used in metastatic colorectal cancer [21]. It has been repurposed as a competitive inhibitor of NSP15 by binding to the enzyme’s active site, thereby inhibiting its NendoU activity. However, tipiracil alone is insufficient for blocking viral replication and proliferation. Thus, it would need to be combined with other drugs to inhibit viral replication successfully [21].

Secondary functions of viral proteins can also be potential targets for therapeutics to better combat viral infections. For example, if the catalytic domain of NSP15 is responsible for disrupting the keratin cytoskeleton, then specific drugs could be repurposed to block its catalytic site.

## Conclusions

NSP15 was found to interact with the cytoskeletal keratin via LC-MS and subsequently validated via immunoprecipitation and immunoblot. As a result of this interaction, a disassembly of the keratin cytoskeleton was observed, leading to a reduction in cell viability. The novel protein-protein interaction identified in this study reveals a previously uncharacterized secondary function of NSP15. The knowledge gained in this study could provide new avenues for treating COVID-19 by inhibiting this enzyme’s activity.

## Author Contributions

D.T., C.M-A., H.A., and H.B. were involved in the experimental section and drafted and edited the manuscript. H.B. conceived the study and edited the manuscript. All authors critically reviewed the manuscript. All authors have read and agreed to the published version of the manuscript.

## References

1. Oxford JS. Influenza A pandemics of the 20th century with special reference to 1918: virology, pathology and epidemiology. Rev Med Virol. 2000;10: 119–133. doi:10.1002/(SICI)1099-1654(200003/04)10:2%3C119::AID-RMV272%3E3.0.CO;2-O

2. Brant AC, Tian W, Majerciak V, Yang W, Zheng Z-M. SARS-CoV-2: from its discovery to genome structure, transcription, and replication. Cell Biosci. 2021;11: 136. doi:10.1186/s13578-021-00643-z

3. Yadav R, Chaudhary JK, Jain N, Chaudhary PK, Khanra S, Dhamija P, et al. Role of structural and non-structural proteins and therapeutic targets of SARS-CoV-2 for COVID-19. Cells. 2021;10: 82. doi:10.3390/cells10040821

4. Low ZY, Zabidi NZ, Yip AJW, Puniyamurti A, Chow VTK, Lal SK. SARS-CoV-2 non-structural proteins and their roles in host immune evasion. Viruses. 2022;14: 1991. doi:10.3390/v14091991

5. Redondo N, Zaldívar-López S, Garrido JJ, Montoya M. SARS-CoV-2 accessory proteins in viral pathogenesis: Knowns and unknowns. Front Immunol. 2021;12. doi:10.3389/fimmu.2021.708264

6. Thoms M, Buschauer R, Ameismeier M, Koepke L, Denk T, Hirschenberger M, et al. Structural basis for translational shutdown and immune evasion by the Nsp1 protein of SARS-CoV-2. Science. 2020;369: 1249–1255. doi:10.1126/science.abc8665

7. Lei X, Dong X, Ma R, Wang W, Xiao X, Tian Z, et al. Activation and evasion of type I interferon responses by SARS-CoV-2. Nat Commun. 2020;11: 3810. doi:10.1038/s41467-020-17665-9

8. Li W, Qiao J, You Q, Zong S, Peng Q, Liu Y, et al. SARS-CoV-2 Nsp5 Activates NF-κB Pathway by Upregulating SUMOylation of MAVS. Front Immunol. 2021;12. doi:10.3389/fimmu.2021.750969

9. Shemesh M, Aktepe TE, Deerain JM, McAuley JL, Audsley MD, David CT, et al. SARS-CoV-2 suppresses IFNβ production mediated by NSP1, 5, 6, 15, ORF6 and ORF7b but does not suppress the effects of added interferon. PLOS Pathog. 2021;17: e1009800. doi:10.1371/journal.ppat.1009800

10. Pillon MC, Frazier MN, Dillard LB, Williams JG, Kocaman S, Krahn JM, et al. Cryo-EM structures of the SARS-CoV-2 endoribonuclease Nsp15 reveal insight into nuclease specificity and dynamics. Nat Commun. 2021;12: 636. doi:10.1038/s41467-020-20608-z

11. Liu F, Gu J. Retinoic acid inducible gene-I, more than a virus sensor. Protein Cell. 2011;2: 351–357. doi:10.1007/s13238-011-1045-y

12. Bach H, Papavinasasundaram KG, Wong D, Hmama Z, Av-Gay Y. *Mycobacterium tuberculosis* virulence is mediated by PtpA dephosphorylation of human vacuolar protein sorting 33B. Cell Host Microbe. 2008;3: 316–322. doi:10.1016/j.chom.2008.03.008

13. Rohaim MA, El Naggar RF, Clayton E, Munir M. Structural and functional insights into non-structural proteins of coronaviruses. Microb Pathog. 2021;150: 104641. doi:10.1016/j.micpath.2020.104641

14. Moll R, Franke WW, Volc-Platzer B, Krepler R. Different keratin polypeptides in epidermis and other epithelia of human skin: a specific cytokeratin of molecular weight 46,000 in epithelia of the pilosebaceous tract and basal cell epitheliomas. J Cell Biol. 1982;95: 285–295. doi:10.1083/jcb.95.1.285

15. Goodson HV, Jonasson EM. Microtubules and microtubule-associated proteins. Cold Spring Harb Perspect Biol. 2018;10: a022608. doi:10.1101/cshperspect.a022608

16. Roberts S, Ashmole I, Rookes SM, Gallimore PH. Mutational analysis of the human papillomavirus type 16 E1-E4 protein shows that the C terminus is dispensable for keratin cytoskeleton association but is involved in inducing disruption of the keratin filaments. J Virol. 1997;71: 3554–3562. doi:10.1128/JVI.71.5.3554-3562.1997

17. Murata T, Goshima F, Nishizawa Y, Daikoku T, Takakuwa H, Ohtsuka K, et al. Phosphorylation of cytokeratin 17 by herpes simplex virus type 2 US3 protein kinase. Microbiol Immunol. 2002;46: 707–719. doi:10.1111/j.1348-0421.2002.tb02755.x

18. De Conto F, Conversano F, Razin SV, Belletti S, Arcangeletti MC, Chezzi C, et al. Host-cell dependent role of phosphorylated keratin 8 during influenza A/NWS/33 virus (H1N1) infection in mammalian cells. Virus Res. 2021;295: 198333. doi:10.1016/j.virusres.2021.198333

19. Tam D, Lorenzo-Leal AC, Hernández LR, Bach H. Targeting SARS-CoV-2 non-structural proteins. Int J Mol Sci. 2023;24: 13002. doi:10.3390/ijms241613002

20. Hong S, Seo SH, Woo S-J, Kwon Y, Song M, Ha N-C. Epigallocatechin gallate inhibits the uridylate-specific endoribonuclease Nsp15 and efficiently neutralizes the SARS-CoV-2 strain. J Agric Food Chem. 2021;69: 5948–5954. doi:10.1021/acs.jafc.1c02050

21. Kim Y, Wower J, Maltseva N, Chang C, Jedrzejczak R, Wilamowski M, et al. Tipiracil binds to uridine site and inhibits Nsp15 endoribonuclease NendoU from SARS-CoV-2. Commun Biol. 2021;4: 193. doi:10.1038/s42003-021-01735-9

